# Docking and SPR identify candidate interactions between Apolipoprotein A-I and chemokine receptors

**DOI:** 10.64898/2026.11.14.766677

**Authors:** Caron Romel, Xavier Blanchet, Sanne L. Maas, Emiel P.C. van der Vorst

**Author notes:** Denotes shared last-authorship (Sanne L. Maas, Emiel P.C. van der Vorst). **Correspondence:** Emiel van der Vorst, University Hospital Aachen, Pauwelsstrasse 30, Aachen, Germany.

## Abstract

**Background:** Apolipoprotein A1 (ApoA1), the main protein component of high-density lipoprotein, suppresses chemokine-driven leukocyte migration. This effect is usually attributed to indirect mechanisms such as cholesterol efflux, changes in lipid rafts, and altered receptor expression. Whether ApoA1 interacts with chemokine receptors directly is not known.

**Methods:** We combined *in silico* rigid-body docking of AlphaFold-predicted structures (GRAMM-X, with interface analysis by PDBePISA) and surface plasmon resonance (SPR). In SPR experiments, ApoA1 was immobilized on the sensor, and CCR1, CCR5, CX_3_CR1, CXCR2, and CXCR4 were incorporated into proteoliposomes and injected as analytes. Mock proteoliposomes lacking any receptor served as a negative control. Kinetics were fitted with a 1:1 Langmuir model.

**Results:** Docking predicted favorable interfaces between ApoA1 and all five receptors. CCR1 had the most favorable docking interaction, and CCR5 had the largest predicted interface (3716.7 Å²; ΔiG −62.8 kcal/mol). Mock proteoliposomes showed no detectable binding to ApoA1, whereas all five receptors bound to ApoA1 in the SPR experiments. The apparent equilibrium dissociation constants (K_D_) were 3.44×10□^8^ M for CXCR4, 3.96×10□^8^ M for CXCR2, 5.26×10□^8^ M for CCR5, 4.35×10□^7^ M for CCR1, and 1.23×10□^6^ M for CX_3_CR1.

**Conclusions:** ApoA1 appears to bind chemokine receptors directly, with the highest apparent affinities for CXCR4, CXCR2, and CCR5. These findings suggest a candidate mechanism, complementary to cholesterol efflux and receptor expression regulation, by which ApoA1 may influence chemokine signaling. However, whether binding alters receptor signaling, internalization, or cell migration remains to be determined.

## Introduction

Apolipoprotein A1 (ApoA1) is the main protein component of high-density lipoprotein (HDL) and an important participant in lipid metabolism^1^. It is synthesized mainly in the liver and intestine, where it helps form and stabilize HDL particles. One of its best-characterized functions is its role in the initial steps of reverse cholesterol transport, in which lipid-poor ApoA1 and small HDL particles promote cholesterol efflux from peripheral cells. The body then processes and transports cholesterol through pathways that can ultimately deliver it to the liver for reuse or excretion. Although HDL cholesterol (HDL-C) levels are inversely associated with atherosclerotic cardiovascular disease^2^, this association does not necessarily indicate that HDL-C itself has a causal protective effect. Moreover, pharmacological strategies that simply raise HDL-C have generally failed to improve cardiovascular outcomes. Consequently, attention has shifted to the functional properties of HDL and ApoA1 rather than plasma HDL-C concentration alone^2^.

Beyond promoting cholesterol efflux, ApoA1 and HDL have been associated with antioxidant, antithrombotic, and anti-inflammatory effects in cardiovascular disease, cancer, and diabetes mellitus^3^. Their anti-inflammatory properties are particularly relevant to innate immune cells, including neutrophils and monocytes/macrophages, and cannot be explained by cholesterol efflux alone^4^. Early work showed that ApoA1 reduced neutrophil degranulation and superoxide production. This study identified saturable ApoA1 binding sites on neutrophils, suggesting a specific interaction that could be a ligand-receptor interaction^5^. Later studies showed that ApoA1 also affects leukocyte migration. Acute exposure of macrophages to human ApoA1 reduced chemotaxis towards a range of chemoattractants, while pre-treatment of monocytes with ApoA1 reduced their recruitment in a murine peritonitis model^6^. These observations indicate that ApoA1 can suppress cell migration, but do not establish the underlying mechanism.

One proposed indirect mechanism involves ApoA1-mediated cholesterol efflux through ABCA1 and ABCG1. Changes in membrane cholesterol content and lipid-raft organization can, in turn, alter receptor localization and signaling. Such effects have been reported for the IL-3/GM-CSF receptor β subunit in hematopoietic stem cells and for TLR4 in macrophages^7,8^. ApoA1 has also been reported to reduce the expression of chemokine receptors, including CCR2 and CX_3_CR1, and of their ligands CCL2, CCL5 and CX_3_CL1^9^. However, the possibility that ApoA1 binds directly to chemokine receptors has received little systematic attention. Because ApoA1 has been reported to affect migration in response to multiple chemoattractants, it is also worth asking whether it acts on a shared component of chemotactic signaling. These observations motivate investigation of direct receptor interactions, but do not themselves establish that such interactions occur.

Chemokine receptors are plausible candidate binding partners. These G protein-coupled receptors (GPCRs) control leukocyte trafficking and are tightly regulated by ligand availability, desensitization and internalization, oligomerization, and their membrane environment, including membrane lipid composition^10^. Several family members, including CCR5, CXCR4 and CXCR2, are important therapeutic targets in cancer, inflammation and viral infection^11,12^. A direct interaction would provide an additional route by which an abundant circulating protein could modulate chemokine signaling and might help explain how ApoA1 affects the migration of cells responding to different chemokines.

Here, we asked whether ApoA1 can bind directly to chemokine receptors. We combined *in silico* docking with real-time binding measurements to investigate five receptors, CCR1, CCR5, CX_3_CR1, CXCR2, and CXCR4, selected for their distinct ligand specificities, immune-cell expression patterns, and roles in inflammatory disease.

## Materials and Methods

### Protein docking

Rigid-body protein-protein docking was performed using the GRAMM-X protein-protein docking web-based platform (v1.2.0)^13^. The full protein structures of ApoA1, CCR1, CCR5, CX_3_CR1, CXCR2, and CXCR4 were obtained from the AlphaFold Protein Structure Database (v6)^14^. For each protein-protein interaction, we predicted 10 docking models. The obtained models were visualized using PyMOL (v3)^15^, and the predicted protein-protein interfaces were further analyzed using the Proteins, Interfaces, Structures and Assemblies (PDBePISA) web-based tool^16^ to calculate interface areas and solvation free energy (Δ^i^G). According to the PDBePISA documentation, *P*-values below 0.5 indicate interfaces with unexpectedly high hydrophobicity relative to randomly selected surface patches, suggesting potential interaction specificity.

### Surface plasmon resonance

Surface plasmon resonance (SPR) experiments were performed using a Sierra SPR®-24 Pro instrument (Bruker) equipped with an amine flat sensor (Bruker). For ligand immobilization, 1 µg/mL of ApoA1 (Bio-rad) in sodium acetate (pH 4.5) was immobilized at a flow rate of 10 µL/min for 360 sec on the sensor surface via amine coupling, resulting in an average immobilization level of 480.4 resonance units (RU). The running buffer contained 10 mM HEPES, 150 mM NaCl, 3 mM EDTA, and 0.005% Tween 20. The wash buffer was Milli-Q water containing 0.05% Tween 20.

The chemokine receptors CCR1, CCR5, CX_3_CR1, CXCR2, and CXCR4 were encapsulated in proteoliposomes (All Abnova) and injected as analytes. Mock proteoliposomes (Abnova) lacking the receptor were included as a negative control. The analytes were injected at five concentrations in a twofold serial dilution series, ranging from 0.125 µg/mL to 2 µg/mL at a flow rate of 60 µL/min, with a 90 sec association phase followed by a 120 sec dissociation phase. The sensor surface was regenerated using 10 mM glycine at pH 2.0.

Sensorgrams were analyzed using Bruker SPR Analyzer 4 software (v4.4.0). Binding kinetics were fitted using a 1:1 Langmuir binding model to determine the association and dissociation rate constants and the kinetic K_D_.

## Results

### *In silico* docking predicts candidate interactions between chemokine receptors and ApoA1

To assess whether ApoA1 could interact directly with chemokine receptors, we first modeled the interactions *in silico*. In the absence of experimental structures of ApoA1-receptor complexes, we used AlphaFold-predicted structures of human ApoA1 and of CCR1, CCR5, CX_3_CR1, CXCR2 and CXCR4. ApoA1 was docked to each receptor with the rigid-body server GRAMM-X, which generated 10 models per protein pair. We analyzed the predicted interfaces with PDBePISA and compared interface area and the solvation free energy (ΔiG). These results were used to rank the models. Because rigid-body docking does not account for conformational change and its energy estimates are approximate, we treated these predicted poses as hypotheses for experimental testing.

The most favourable binding was analysed based on the lowest *P*-value of solvation free energy, followed by interface area. The selected docking model for each receptor is shown in **Figure 1A-E**, the other nine models are presented in **Supplementary Figures 1-5**. Among the receptors analyzed, CCR1 showed the most favorable docking interaction, with the lowest *P*-value of 0.068, an interface area of 2295.9 Å², and a predicted Δ^i^G of -41.8 kcal/mol. However, CCR5 showed the largest predicted receptor-ApoA1 interface area, with an interface area of 3716.7 Å² and a Δ^i^G of -62.8 kcal/mol. CX_3_CR1, CXCR2, and CXCR4 also demonstrated favorable docking interactions with ApoA1, with all models showing potential hydrophobic contacts. Thereby, these results clearly predict that there are preferential binding sites between ApoA1 and different chemokine receptors.

**Figure 1.**
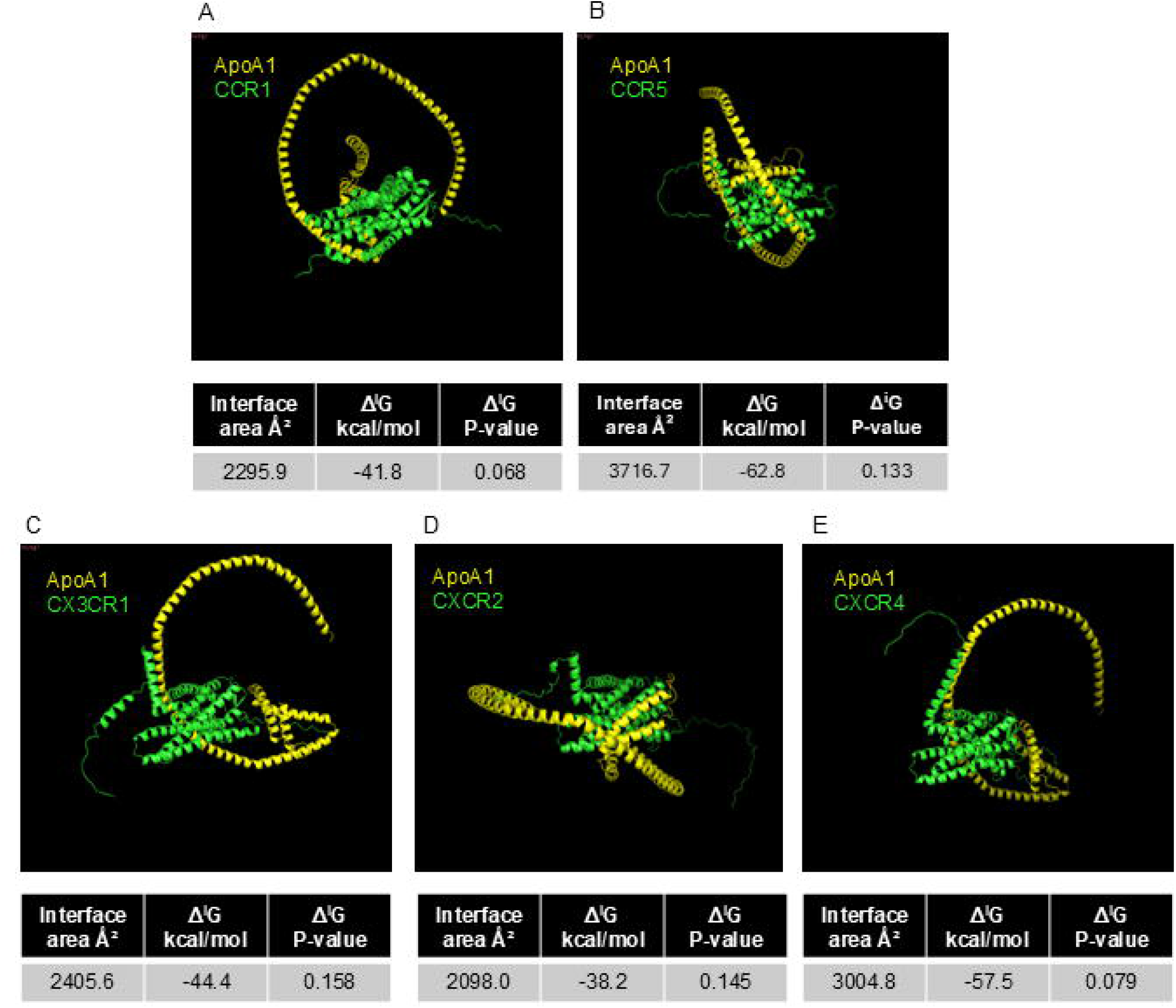
ApoA1-chemokine receptor docking models and interface analysis. Representative images show the selected docking models of ApoA1 and (A) CCR1, (B) CCR5, (C) CX_3_CR1, (D) CXCR2, and (E) CXCR4, which were generated using GRAMM-X. The interface area, solvation free-energy gain (Δ^i^G), and Δ^i^G *P*-value for each predicted interface were calculated using PDBePISA. ApoA1 - Apolipoprotein A1, CCR - CC Chemokine Receptors, CX_3_CR1 - CX3C motif chemokine receptor 1, CXCR - C-X-C motif chemokine receptor, k_a_ - Association rate constant, k_d_ - dissociation rate constant, K_D_ - equilibrium dissociation constant, PDBePISA - Proteins, Interfaces, Structures and Assemblies.

### SPR validation of ApoA1 binding to chemokine receptors

To test whether ApoA1 binds the receptors experimentally, we measured real-time binding kinetics between ApoA1 and the respective chemokine receptors using SPR. For the binding experiments, the receptors were incorporated into proteoliposomes to maintain structural integrity and provide a membrane-like environment. Mock proteoliposomes lacking any receptor showed no detectable binding response to ApoA1 under the tested conditions, indicating that the proteoliposome carrier itself does not bind ApoA1 (**Figure 2A**). Binding responses were detected for proteoliposomes containing each of the five chemokine receptors (**Figure 2B-F**). The sensorgrams were analyzed using a 1:1 Langmuir binding model.

**Figure 2.**
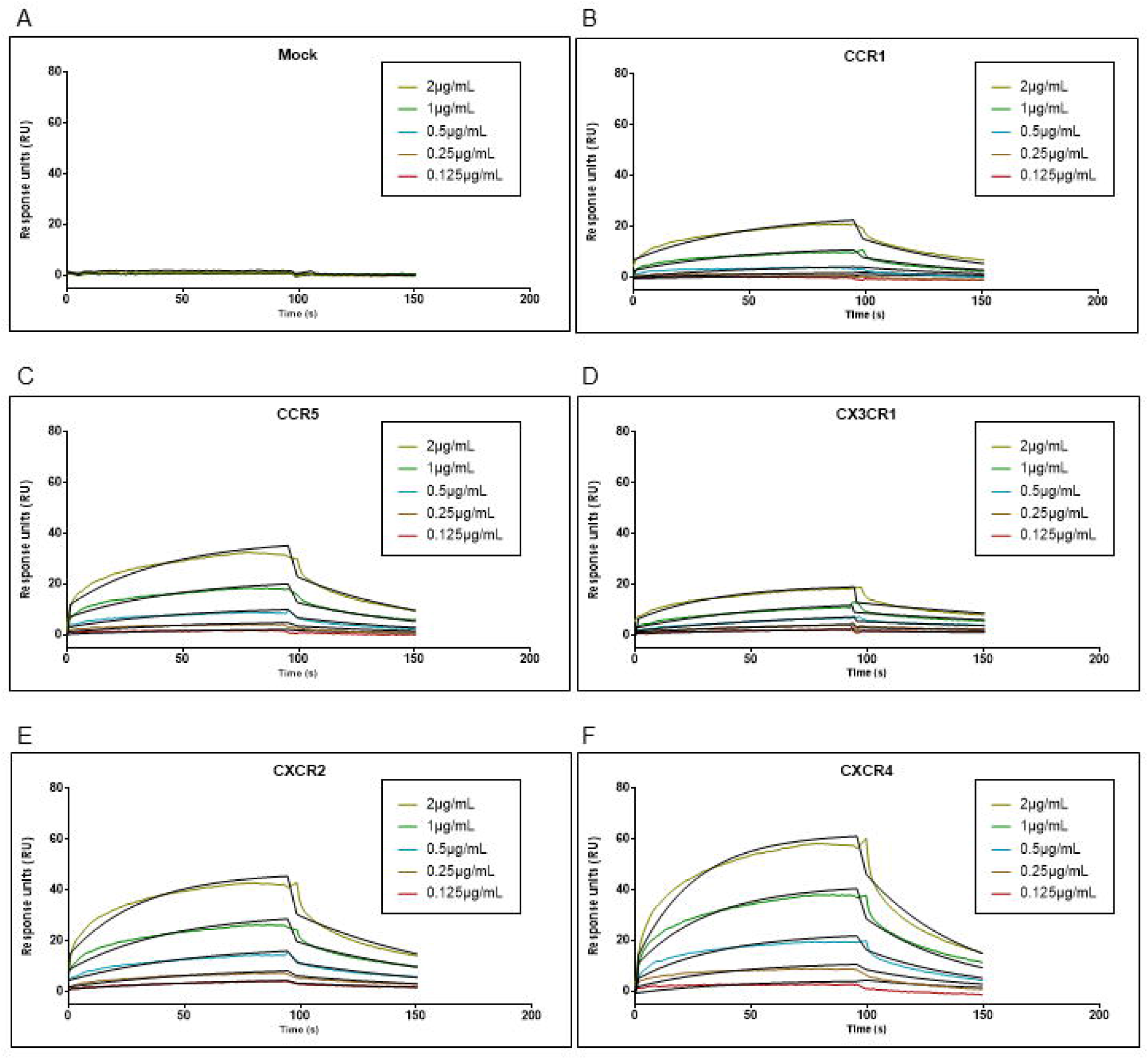
SPR sensorgrams showing the binding responses of ApoA1 to chemokine receptors-containing proteoliposomes. SPR sensorgrams for ApoA1 binding to (A) mock proteoliposomes, and proteoliposomes containing (B) CCR1, (C) CCR5, (D) CX_3_CR1, (E) CXCR2, and (F) CXCR4. Each analyte was tested at five concentrations: 2, 1, 0.5, 0.25, and 0.125 µg/mL. Binding kinetics were analysed using a 1:1 Langmuir binding model. ApoA1 - Apolipoprotein A1, CCR - CC Chemokine Receptors, CX_3_CR1 - CX3C motif chemokine receptor 1, CXCR - C-X-C motif chemokine receptor, SPR - Surface plasmon resonance.

The SPR analysis demonstrated distinct binding kinetics for ApoA1 binding with all five chemokine receptors. Although the experimentally determined binding affinities did not follow the same order as the predicted interactions from the *in-silico* docking analysis, all five receptors showed measurable binding responses to ApoA1. CXCR4 showed the lowest K_D_ (3.44×10□^8^ M), followed by CXCR2 (3.96×10□^8^ M) and CCR5 (5.26×10□^8^ M). CCR1 and CX_3_CR1 showed higher K_D_ values of 4.35×10□^7^ M and 1.23×10□^6^ M, respectively. **Table 1** summarizes the corresponding association and dissociation rate constants for each receptor. Altogether, these SPR validation experiments indicate direct binding between ApoA1 and several chemokine receptors.

**Table 1.**
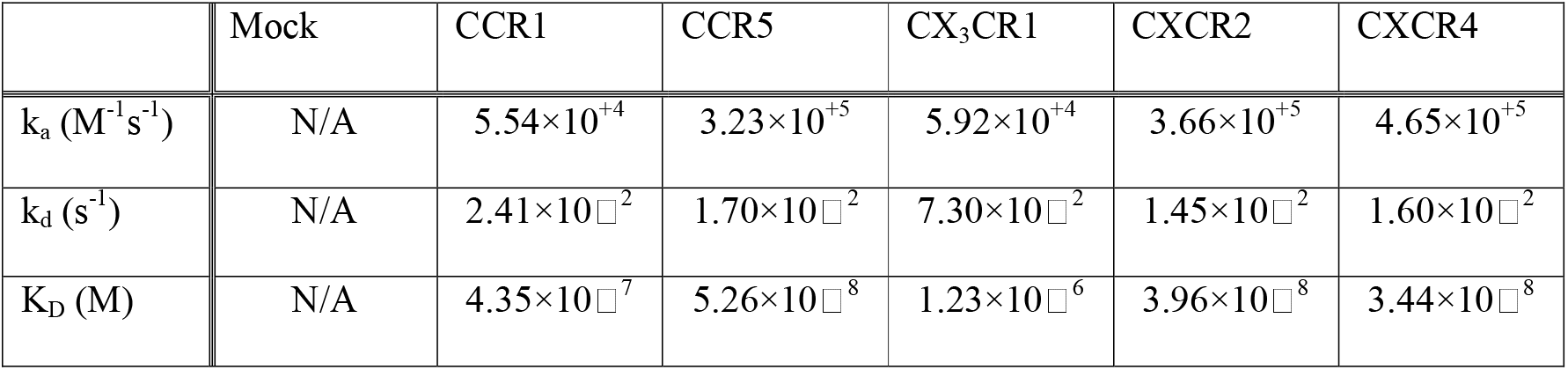
SPR-derived binding kinetics of ApoA1 and chemokine receptors-containing proteoliposomes. Association rate constant (k_a_), dissociation rate constant (k_d_), and equilibrium dissociation constant (K_D_) were obtained by fitting the SPR data using a 1:1 Langmuir binding model. N/A: not available. ApoA1 - Apolipoprotein A1, CCR - CC Chemokine Receptors, CX_3_CR1 - CX3C motif chemokine receptor 1, CXCR - C-X-C motif chemokine receptor 2, SPR - Surface plasmon resonance

|  | Mock | CCR1 | CCR5 | CX<sub>3</sub>CR1 | CXCR2 | CXCR4 |
|---|---|---|---|---|---|---|
| k<sub>a</sub> (M<sup>-1</sup>s<sup>-1</sup>) | N/A | 5.54×10<sup>+4</sup> | 3.23×10<sup>+5</sup> | 5.92×10<sup>+4</sup> | 3.66×10<sup>+5</sup> | 4.65×10<sup>+5</sup> |
| k<sub>d</sub> (s<sup>-1</sup>) | N/A | 2.41×10□<sup>2</sup> | 1.70×10□<sup>2</sup> | 7.30×10□<sup>2</sup> | 1.45×10□<sup>2</sup> | 1.60×10□<sup>2</sup> |
| K<sub>D</sub> (M) | N/A | 4.35×10□<sup>7</sup> | 5.26×10□<sup>8</sup> | 1.23×10□<sup>6</sup> | 3.96×10□<sup>8</sup> | 3.44×10□<sup>8</sup> |

## Discussion

In this study, ApoA1 bound all five chemokine receptor-containing proteoliposomes tested (CCR1, CCR5, CX_3_CR1, CXCR2 and CXCR4) in an SPR assay. Mock proteoliposomes showed no detectable binding under the tested conditions, suggesting the observed responses were due to the presence of a receptor in the proteoliposomes rather than the mock lipid carrier alone. These results suggest that ApoA1 may physically interact with chemokine receptors, a mechanism not previously considered as a potential explanation for ApoA1’s effects on leukocyte migration.

The docking metrics and SPR-derived affinity rankings were not fully concordant. CCR5 had the largest predicted interface area and most favorable ΔiG, whereas CXCR4 showed the lowest apparent K_D_ by SPR. Although CCR1 had the lowest reported ΔiG *P*-value, its apparent SPR affinity was lower than that of CCR5, with an approximately 8.3-fold higher K_D_. Several features of the approach may explain this discrepancy. Rigid-body docking treats both partners as fixed, whereas GPCRs and ApoA1 are conformationally flexible. AlphaFold models of GPCRs are predicted without a membrane or ligand, and interface area and ΔiG from docking do not reliably predict binding affinitiy^17,18^. We therefore regard the docking results as a way to identify candidate interaction surfaces, rather than to predict affinity. The models may guide mutagenesis of candidate interface residues to test whether the predicted contacts contribute to binding.

The affinity pattern may have biological implications, as the three tightest binders, CXCR4, CXCR2, and CCR5, are major regulators of neutrophil, monocyte, and lymphocyte trafficking and are therapeutic targets in inflammation, cancer, and viral infection^10–12^. CXCR2 is of particular interest because ApoA1 and ApoA1 mimetic peptides have been reported to suppress CXCR2-directed neutrophil chemotaxis and reduce CXCR2 surface expression on neutrophils^19^. Direct association with CXCR2 could contribute to these effects, for example by changing receptor availability or internalization, although our study did not test this possibility. ApoA1 is present in the plasma at micromolar concentrations, mostly in HDL. The affinities measured here are therefore within a physiologically plausible concentration range. However, differences in ApoA1 presentation and the assay format mean that this comparison does not establish physiological relevance.

Our results complement an earlier study showing that ApoA1 reduced the expression of CCR2, CX_3_CR1, and their ligands in monocytes and endothelial cells through NF-κB and PPARγ signalling^9^. Their proposed mechanism involves gene regulation, whereas ours provides evidence consistent with physical association with receptor-containing proteoliposomes. These mechanisms could coexist. CX_3_CR1 is informative in this respect. It was the weakest binder in our assay, yet ApoA1 has been reported to regulate its expression^9^. This suggests that binding affinity and gene-regulatory effects may be partly independent, and that a given receptor can be affected by one mechanism, the other, or both.

Several mechanisms could follow from a direct interaction, but our data do not distinguish between them. ApoA1 might compete with chemokines for binding, or it might bind at a different site and act as an allosteric modulator. It could also promote receptor clustering or internalization or alter how the receptor sits in the membrane. Because ApoA1 is a lipid-binding protein, it may also influence the lipid environment near the receptor. Distinguishing among these possibilities will require functional experiments.

Our study has limitations. First, the SPR format has caveats. ApoA1 was immobilized by amine coupling, which produces random orientation and may mask binding sites. The analytes were proteoliposomes, so receptor density and orientation are not controlled, and multivalent binding can inflate apparent affinity. Because the three highest-affinity receptors differ by less than 1.5-fold, their ranking should not be over-interpreted. Second, we measured binding responses but did not test function. Whether the interaction alters signaling, internalization, or cell migration is unknown and warrants future investigation.

In summary, our data support an association between ApoA1 and proteoliposomes containing chemokine receptors, with the strongest apparent affinities for CXCR4, CXCR2, and CCR5. This adds a candidate mechanism to existing explanations of how ApoA1 affects the chemokine system and provides a framework for testing the functional consequences of these interactions.

## Data availability

All data are contained within the manuscript.

## Disclosure statement

All the authors declared no competing interests.

## Funding

This work was supported by the Corona Foundation (S199/10084/2021), and by the Deutsche Forschungsgemeinschaft (DFG) (SFB TRR219 - Project-ID 322900939; subproject M07, Project-ID 569469700 and INST 222/1408-1 FUGG; Projekt: 460284161) to E.P.C.v.d.V.

## Author contributions

Conceptualization: E.P.C.v.d.V.

Data curation: C.R., X.B.

Formal analysis: C.R., X.B.

Supervision: S.L.M., E.P.C.v.d.V.

Writing – Original draft: C.R.

Writing – review and editing: S.L.M., E.P.C.v.d.V.

## Notes

### Competing Interest Statement

The authors have declared no competing interest.

