## Supplemental Figures for "Docking and SPR identify candidate interactions between Apolipoprotein A-I and chemokine receptors"

*Denotes shared last-authorship

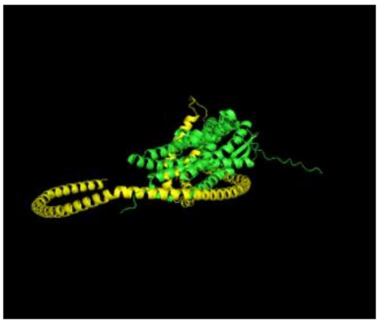

| Interface area Å² | $\Delta^i G$ kcal/mol | $\Delta^i G$ P-value |
|---|---|---|
| 2236.1 | -38.6 | 0.214 |

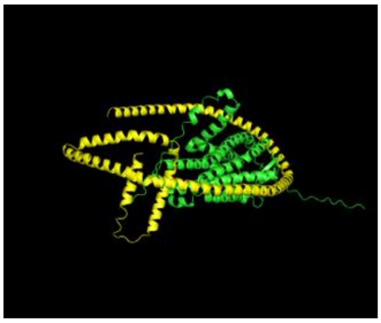

| Interface area Å² | $\Delta^i G$ kcal/mol | $\Delta^i G$ P-value |
|---|---|---|
| 3125.1 | -43.0 | 0.453 |

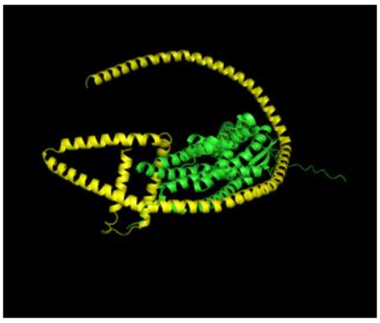

| Interface area Å² | $\Delta^i G$ kcal/mol | $\Delta^i G$ P-value |
|---|---|---|
| 2714.5 | -39.0 | 0.409 |

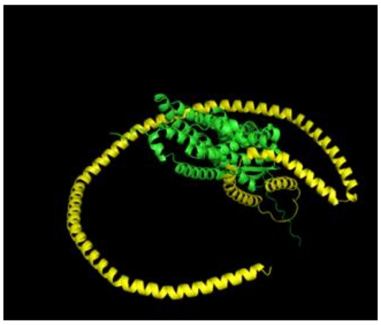

| Interface area Å² | $\Delta^i G$ kcal/mol | $\Delta^i G$ P-value |
|---|---|---|
| 2403.1 | -43.9 | 0.160 |

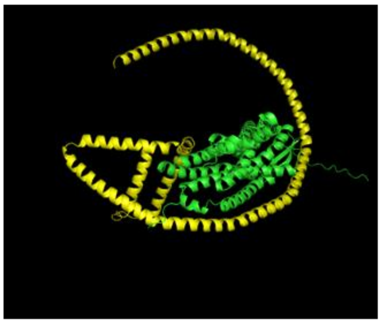

| Interface area Å² | $\Delta^i G$ kcal/mol | $\Delta^i G$ P-value |
|---|---|---|
| 2480.6 | -38.6 | 0.336 |

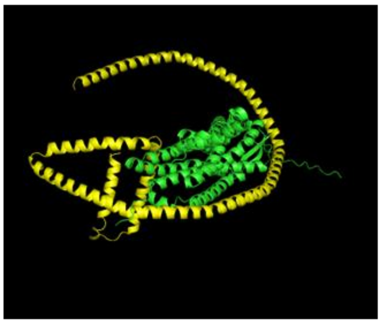

| Interface area Å² | $\Delta^i G$ kcal/mol | $\Delta^i G$ P-value |
|---|---|---|
| 2705.5 | -36.0 | 0.486 |

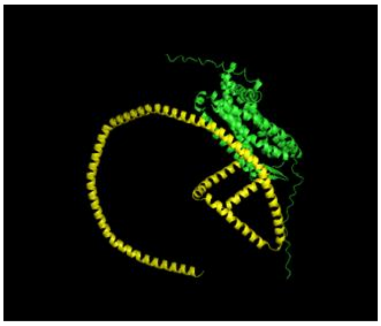

| Interface area Å² | $\Delta^i G$ kcal/mol | $\Delta^i G$ P-value |
|---|---|---|
| 1930.2 | -26.9 | 0.477 |

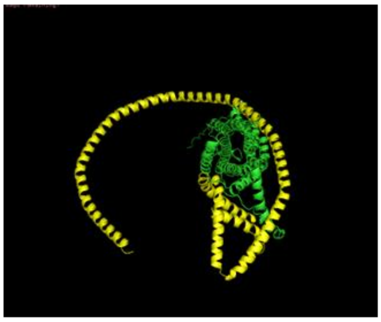

| Interface area Å² | $\Delta^i G$ kcal/mol | $\Delta^i G$ P-value |
|---|---|---|
| 2663.3 | -37.3 | 0.393 |

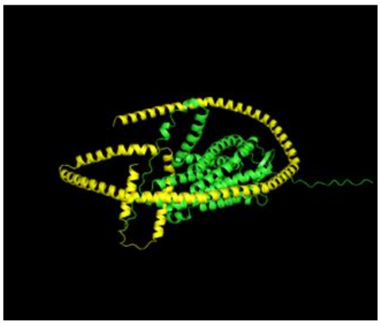

| Interface area Å² | $\Delta^i G$ kcal/mol | $\Delta^i G$ P-value |
|---|---|---|
| 2288.7 | -32.9 | 0.504 |

**Supplementary Figure 1. ApoA1-CCR1 docking models and interface analysis.** The nine additional docking models of ApoA1 (yellow) and CCR1 (green), generated using GRAMM-X. Interface area, solvation free-energy gain ($\Delta^i G$), and $\Delta^i G$ *P*-value for each predicted interface were calculated using PDBePISA. ApoA1 - Apolipoprotein A1, CCR - CC Chemokine Receptors, PDBePISA - Proteins, Interfaces, Structures and Assemblies.

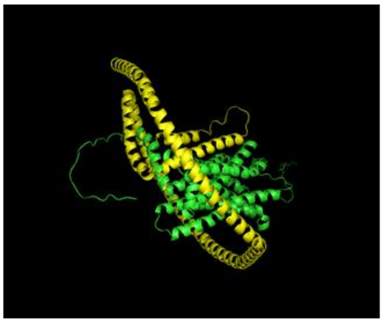

| Interface area Å² | $\Delta^{i}G$ kcal/mol | $\Delta^{i}G$ P-value |
|---|---|---|
| 3434.4 | -62.3 | 0.093 |

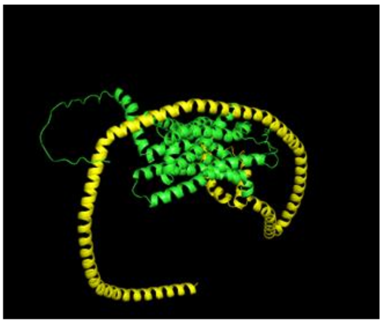

| Interface area Å² | $\Delta^{i}G$ kcal/mol | $\Delta^{i}G$ P-value |
|---|---|---|
| 2760.9 | -47.2 | 0.240 |

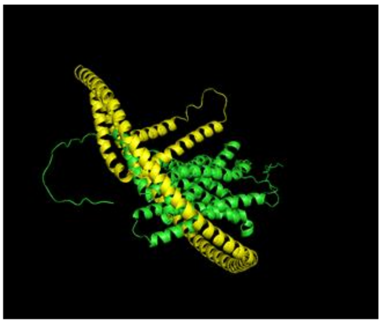

| Interface area Å² | $\Delta^{i}G$ kcal/mol | $\Delta^{i}G$ P-value |
|---|---|---|
| 2944.7 | -53.1 | 0.133 |

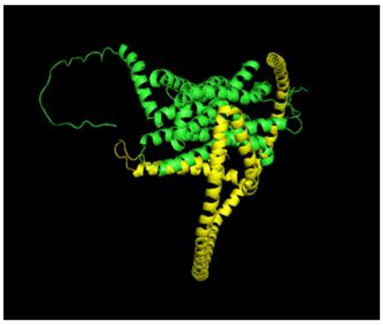

| Interface area Å² | $\Delta^{i}G$ kcal/mol | $\Delta^{i}G$ P-value |
|---|---|---|
| 2333.3 | -34.3 | 0.404 |

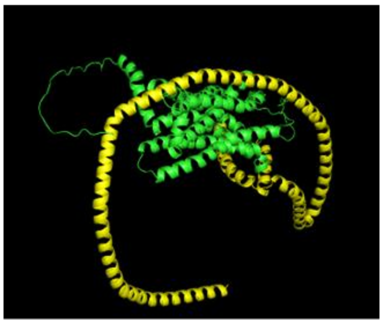

| Interface area Å² | $\Delta^{i}G$ kcal/mol | $\Delta^{i}G$ P-value |
|---|---|---|
| 2422.2 | -43.3 | 0.236 |

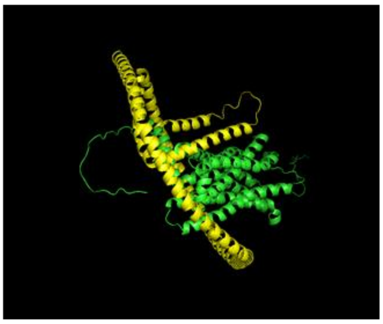

| Interface area Å² | $\Delta^{i}G$ kcal/mol | $\Delta^{i}G$ P-value |
|---|---|---|
| 2771.5 | -46.0 | 0.198 |

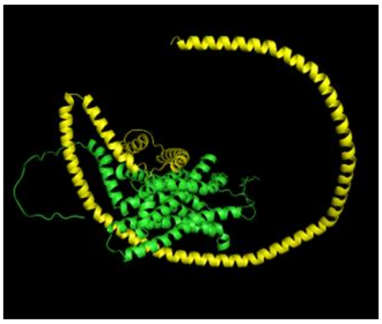

| Interface area Å² | $\Delta^{i}G$ kcal/mol | $\Delta^{i}G$ P-value |
|---|---|---|
| 3552.4 | -51.9 | 0.347 |

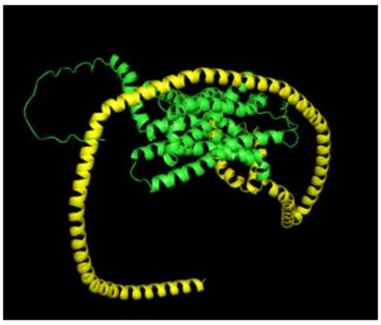

| Interface area Å² | $\Delta^{i}G$ kcal/mol | $\Delta^{i}G$ P-value |
|---|---|---|
| 2738.4 | -35.0 | 0.421 |

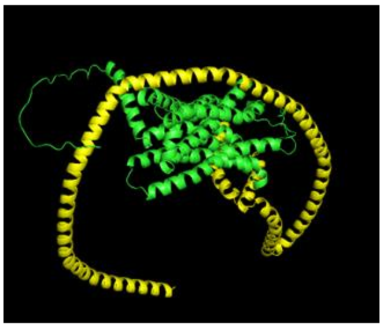

| Interface area Å² | $\Delta^{i}G$ kcal/mol | $\Delta^{i}G$ P-value |
|---|---|---|
| 1878.0 | -22.8 | 0.558 |

**Supplementary Figure 2. Additional ApoA1-CCR5 docking poses.** The nine additional poses of ApoA1 (yellow) and CCR5 (green) were generated using GRAMM-X. For each pose, the interface area, solvation free-energy gain ($\Delta^{i}G$), and $\Delta^{i}G$ *P*-value were obtained with PDBePISA. ApoA1 - Apolipoprotein A1, CCR - CC Chemokine Receptors, PDBePISA - Proteins, Interfaces, Structures and Assemblies.

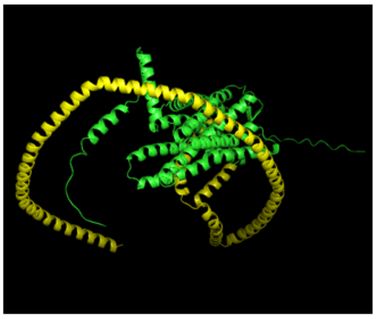

| Interface area Å$^2$ | $\Delta^{i}G$ kcal/mol | $\Delta^{i}G$ P-value |
|---|---|---|
| 2658.6 | -43.8 | 0.330 |

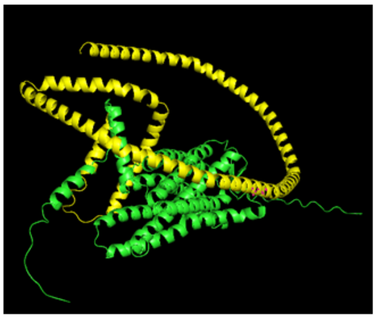

| Interface area Å$^2$ | $\Delta^{i}G$ kcal/mol | $\Delta^{i}G$ P-value |
|---|---|---|
| 2871.7 | -44.3 | 0.338 |

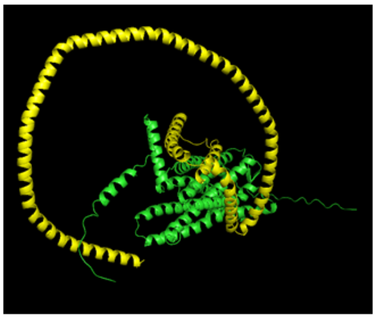

| Interface area Å$^2$ | $\Delta^{i}G$ kcal/mol | $\Delta^{i}G$ P-value |
|---|---|---|
| 2865.0 | -44.2 | 0.415 |

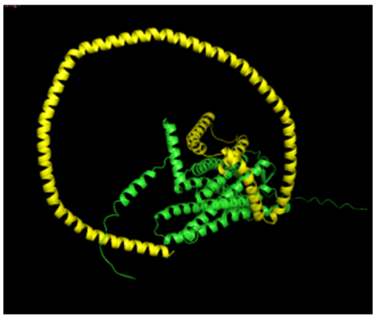

| Interface area Å$^2$ | $\Delta^{i}G$ kcal/mol | $\Delta^{i}G$ P-value |
|---|---|---|
| 2120.7 | -29.4 | 0.392 |

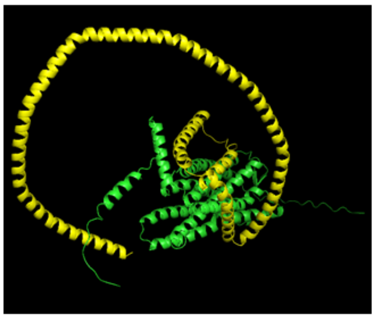

| Interface area Å$^2$ | $\Delta^{i}G$ kcal/mol | $\Delta^{i}G$ P-value |
|---|---|---|
| 2295.0 | -32.7 | 0.495 |

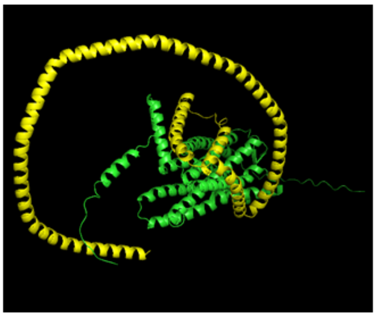

| Interface area Å$^2$ | $\Delta^{i}G$ kcal/mol | $\Delta^{i}G$ P-value |
|---|---|---|
| 2374.8 | -39.7 | 0.349 |

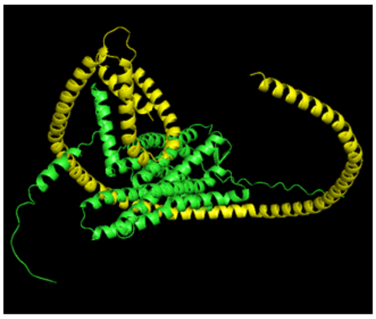

| Interface area Å$^2$ | $\Delta^{i}G$ kcal/mol | $\Delta^{i}G$ P-value |
|---|---|---|
| 2644.4 | -26.6 | 0.848 |

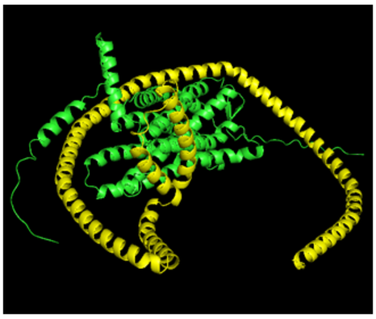

| Interface area Å$^2$ | $\Delta^{i}G$ kcal/mol | $\Delta^{i}G$ P-value |
|---|---|---|
| 2401.7 | -27.9 | 0.715 |

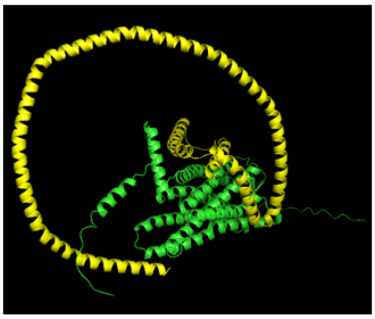

| Interface area Å$^2$ | $\Delta^{i}G$ kcal/mol | $\Delta^{i}G$ P-value |
|---|---|---|
| 2500.4 | -32.1 | 0.598 |

**Supplementary Figure 3. ApoA1-$CX_3CR1$ docking models and interface metrics.** The nine additional docking models, generated by GRAMM-X with ApoA1 (yellow) and $CX_3CR1$ (green) . For each model, interface area, solvation free- energy gain ($\Delta^{i}G$), and *P*-value were calculated using PDBePISA. ApoA1 - Apolipoprotein A1, $CX_3CR1$ - CX3C motif chemokine receptor 1, PDBePISA - Proteins, Interfaces, Structures and Assemblies.

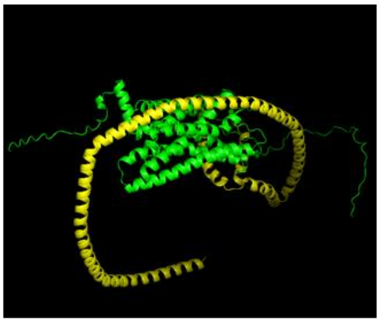

| Interface area Å² | $\Delta^{i}G$ kcal/mol | $\Delta^{i}G$ P-value |
|---|---|---|
| 3032.6 | -42.5 | 0.398 |

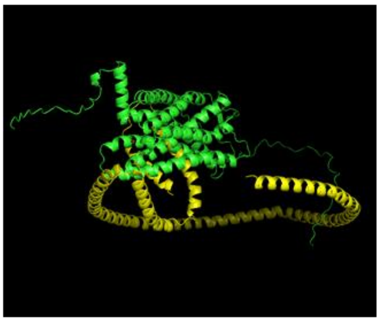

| Interface area Å² | $\Delta^{i}G$ kcal/mol | $\Delta^{i}G$ P-value |
|---|---|---|
| 2900.1 | -48.9 | 0.151 |

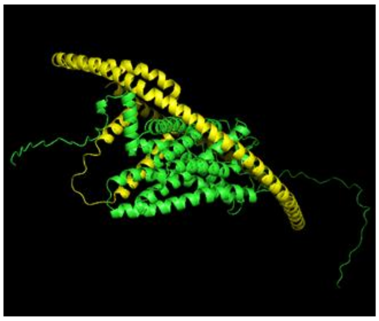

| Interface area Å² | $\Delta^{i}G$ kcal/mol | $\Delta^{i}G$ P-value |
|---|---|---|
| 3047.0 | -52.0 | 0.179 |

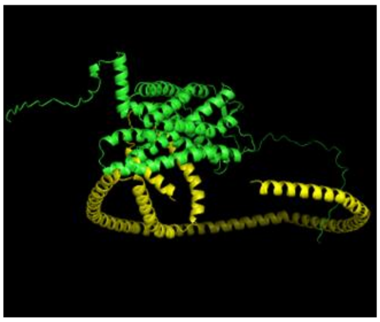

| Interface area Å² | $\Delta^{i}G$ kcal/mol | $\Delta^{i}G$ P-value |
|---|---|---|
| 2006.6 | -32.8 | 0.259 |

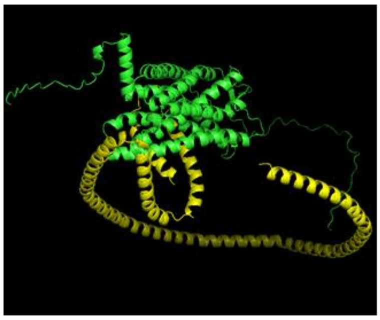

| Interface area Å² | $\Delta^{i}G$ kcal/mol | $\Delta^{i}G$ P-value |
|---|---|---|
| 2125.2 | -39.7 | 0.176 |

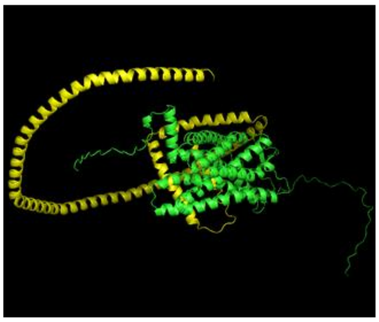

| Interface area Å² | $\Delta^{i}G$ kcal/mol | $\Delta^{i}G$ P-value |
|---|---|---|
| 2441.7 | -44.1 | 0.159 |

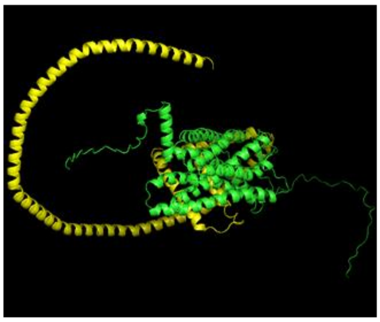

| Interface area Å² | $\Delta^{i}G$ kcal/mol | $\Delta^{i}G$ P-value |
|---|---|---|
| 2463.3 | -38.1 | 0.325 |

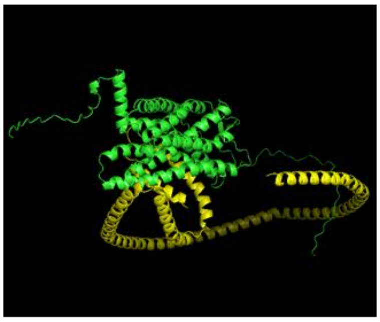

| Interface area Å² | $\Delta^{i}G$ kcal/mol | $\Delta^{i}G$ P-value |
|---|---|---|
| 2229.7 | -36.2 | 0.269 |

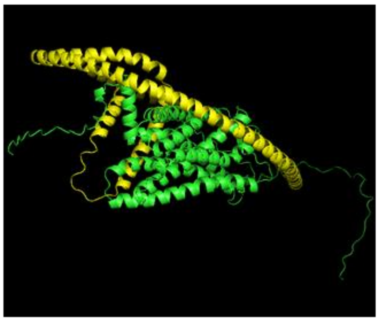

| Interface area Å² | $\Delta^{i}G$ kcal/mol | $\Delta^{i}G$ P-value |
|---|---|---|
| 3350.6 | -50.8 | 0.250 |

**Supplementary Figure 4. ApoA1-CXCR2 poses and interface analysis.** The nine additional poses of ApoA1 (yellow) and CXCR2 (green) were generated using GRAMM-X. The interface area, solvation free- energy gain ($\Delta^{i}G$), and *P*-value for each predicted interface were determined using PDBePISA. ApoA1 - Apolipoprotein A1, CXCR - C-X-C motif chemokine receptor, PDBePISA - Proteins, Interfaces, Structures and Assemblies.

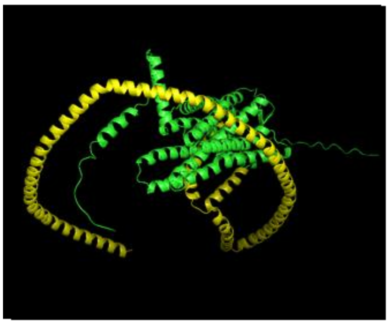

| Interface area Å² | $\Delta^{i}G$ kcal/mol | $\Delta^{i}G$ P-value |
|---|---|---|
| 2687.2 | -53.1 | 0.079 |

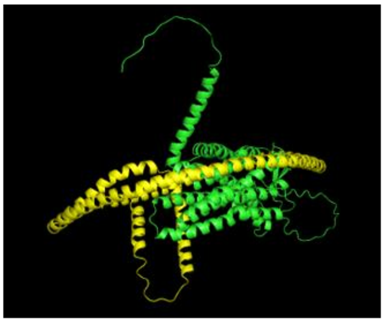

| Interface area Å² | $\Delta^{i}G$ kcal/mol | $\Delta^{i}G$ P-value |
|---|---|---|
| 2169.4 | -34.3 | 0.314 |

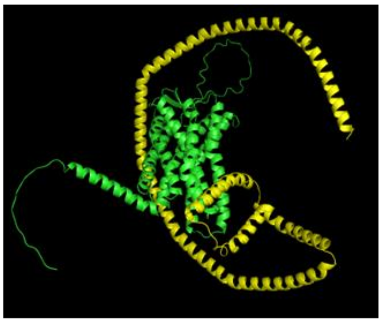

| Interface area Å² | $\Delta^{i}G$ kcal/mol | $\Delta^{i}G$ P-value |
|---|---|---|
| 2078.7 | -35.1 | 0.195 |

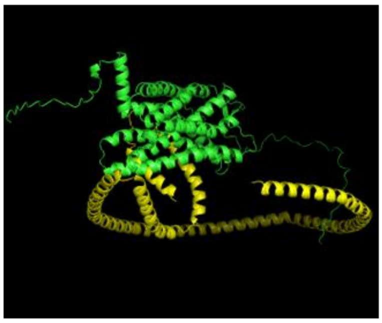

| Interface area Å² | $\Delta^{i}G$ kcal/mol | $\Delta^{i}G$ P-value |
|---|---|---|
| 2290.8 | -37.4 | 0.125 |

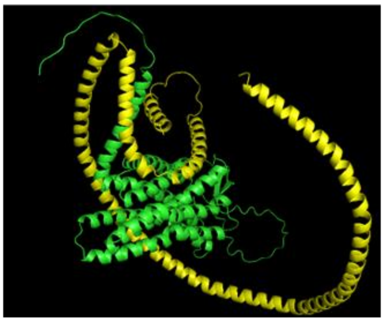

| Interface area Å² | $\Delta^{i}G$ kcal/mol | $\Delta^{i}G$ P-value |
|---|---|---|
| 2917.7 | -42.0 | 0.367 |

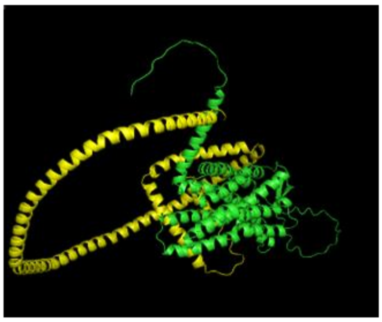

| Interface area Å² | $\Delta^{i}G$ kcal/mol | $\Delta^{i}G$ P-value |
|---|---|---|
| 2515.4 | -44.7 | 0.106 |

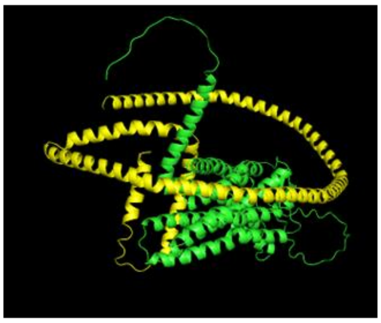

| Interface area Å² | $\Delta^{i}G$ kcal/mol | $\Delta^{i}G$ P-value |
|---|---|---|
| 2846.5 | -40.6 | 0.357 |

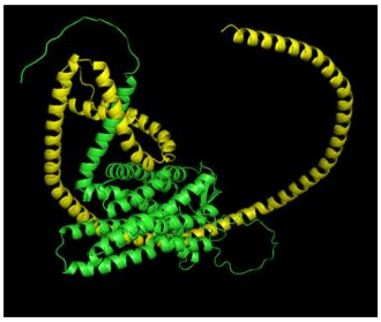

| Interface area Å² | $\Delta^{i}G$ kcal/mol | $\Delta^{i}G$ P-value |
|---|---|---|
| 2692.3 | -27.2 | 0.717 |

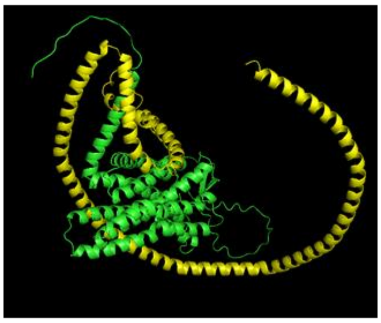

| Interface area Å² | $\Delta^{i}G$ kcal/mol | $\Delta^{i}G$ P-value |
|---|---|---|
| 3364.1 | -38.0 | 0.615 |

**Supplementary Figure 5. ApoA1- CXCR4 docking models and interface analysis.** The nine additional models of ApoA1 (yellow) and CXCR4 (green) were generated using GRAMM-X. The interface area, solvation free- energy gain ($\Delta^{i}G$), and *P*-value were calculated for each predicted interface using PDBePISA. ApoA1 - Apolipoprotein A1, CXCR - C-X-C motif chemokine receptor, PDBePISA - Proteins, Interfaces, Structures and Assemblies.
